# 3D Printed Neural Tissues with *in situ* Optical Dopamine Sensors

**DOI:** 10.1101/2022.07.01.498382

**Authors:** Jianfeng Li, Armin Reimers, Ka My Dang, Michael G. K. Brunk, Jonas Drewes, Ulrike M. Hirsch, Christian Willems, Christian E. H. Schmelzer, Thomas Groth, Ali Shaygan Nia, Xinliang Feng, Fabian Schütt, Wesley D. Sacher, Rainer Adelung, Joyce K. S. Poon

## Abstract

Engineered neural tissues serve as models for studying neurological conditions and drug screening. Besides observing the cellular physiological properties, *in situ* monitoring of neurochemical concentrations with cellular spatial resolution in such neural tissues can provide additional valuable insights in models of disease and drug efficacy. In this work, we demonstrate the first three-dimensional (3D) tissue cultures with embedded optical dopamine (DA) sensors. We developed an alginate/Pluronic F127 based bio-ink for human dopaminergic brain tissue printing with tetrapodal-shaped-ZnO microparticles (t-ZnO) additive as the DA sensor. DA quenches the autofluorescence of t-ZnO in physiological environments, and the reduction of the fluorescence intensity serves as an indicator of the DA concentration. The neurons that were 3D printed with the t-ZnO showed good viability, and extensive 3D neural networks were formed within one week after printing. The t-ZnO can sense DA in the 3D printed neural network with a detection limit of 0.137 μM. The results are a first step toward integrating tissue engineering with intensiometric biosensing for advanced artificial tissue/organ monitoring.

## 1. Introduction

A vast number of debilitating neurological conditions, such as Parkinson’s disease, Alzheimer’s disease, depression, epilepsy and traumatic brain injuries, still do not have effective cures today. Ideally, these conditions are studied in human subjects, *ex vivo* human tissues, or animal models, but the availability of donors and samples, types of samples, and ethics often restrict the studies that can be performed [1, 2]. As a complement to human and animal experiments, neural tissue models, such as *in vitro* two-dimensional (2D) or three-dimensional (3D) tissue cultures and organoids, enable *in situ* investigations of neuronal development with cellular resolution [3]. 3D neuronal tissue models can better recapitulate *in vivo* counterparts than 2D models by emulating physicochemical features in the brain [4]. For 3D neuronal culture, hydrogel, with a high content of water and good biocompatibility, is an ideal platform; it has similar compressive modulus like brain (1.4-3.8 kPa), and enables remodeling during cell migration and differentiation [5]. Many neurological conditions are also strongly affected by neurochemicals (e.g., dopamine (DA), serotonin, and acetylcholine) [6]. Therefore, in neural tissue models, *in situ* measurements of the neurochemical concentrations with cellular spatial resolution can be a powerful functionality for the physiological studies and drug screening.

DA is an important neuromodulator in the central and peripheral nervous systems, regulating movement control, cognition, decision making, and other essential body functions [7]. DA modulates synaptic dynamics and plasticity, affecting the neurotransmitter receptor sensitivity and membrane excitability [8]. Dysfunction of the dopaminergic system can lead to severe nervous system disorders or diseases, such as Parkinson’s disease, schizophrenia and attention deficit hyperactivity disorder [6]. The typical methods to analyze DA are microdialysis and high-performance liquid chromatography (HPLC), which have coarse time and spatial resolution (typically > 1 min and > 100 µm), insufficient to probe cellular activity [9-11]. Fast scan cyclic voltammetry (FSCV) can measure DA with improved temporal (sub-second) and spatial (several microns) resolution [12], but it is limited by sensor drift, microelectrode variation and fragility [13]. Furthermore, microdialysis, HPLC, and FSCV are invasive and may cause tissue damage. Alternatively, optical intensiometric DA sensors provide a non-invasive means to monitor DA concentrations in 3D with high sensitivity [14]. An example is genetically encoded DA optical sensors, such as dLight, which have high selectivity and sensitivity, but transfection is not always possible or effective depending on the cell-type and application [15]. Chemically based DA optical sensors, whose optical absorption, autofluorescence, or other spectral properties change with DA concentration, do not require any genetic modification of the neurons and could thus be more versatile [16]. So far, chemical DA optical sensors, which are often based on carbon-based nanostructures, quantum dots, or silica nanoparticles, have only provided 2D spatial profiles of DA concentration [16, 17].

Here, we present a concept for 3D neural tissue constructs with embedded optical DA sensors. We sought to monitor DA release in 3D with cellular resolution *in situ* by printing the DA sensors and neurons together in 3D scaffolds. We used tetrapodal-shaped-ZnO microparticles (t-ZnO), a biocompatible material ^[18]^ with autofluorescence that could be quenched by DA, as the sensor. Bio-inks formulated with t-ZnO and SH-SY5Y cells were used to 3D print living scaffolds. Prior ZnO-based DA sensors used chiral ZnO nanoparticles and ZnO/carbon dots ^[19, 20]^, but the sensors were 2D, and the nanometer size of the particles may be toxic to living cells [21]. This work shows the potential of 3D neural constructs with integrated optical sensing, paving a way towards new type of smart tissue.

## 2. Results and Discussion

### 2.1. 3D Printing of Neurons with t-ZnO Additive

**Figure 1a** illustrates the overall concept for 3D neural tissue constructs with embedded optical DA sensors. The t-ZnO, which has a ZnO core with four hexagonal arms, and neurons were formulated into a single bio-ink and printed together. Due to the four-armed structure and brittleness, t-ZnO is difficult to be 3D printed and instead often used in powder form or molded into shapes [18]. In our study, Pluronic F127 (Plu) was used as a fugitive matrix to aid t-ZnO 3D printing, which can be removed by sintering to render pure 3D printed t-ZnO constructs. Plu is also widely used for tissue engineering since it is biocompatible and thermally responsive [22, 23]. When printed with living cells, alginate (Alg) was added as a biocompatible cross-linkable material such that the integrity of printed structure can be maintained in an aqueous environment. As will be discussed in Section 2, DA was found to polymerize on the t-ZnO surface in physiological environments, quenching the t-ZnO autofluorescence such that t-ZnO can serve as a DA sensor (**Figure 1b**). For cell laden scaffold printing, cells and t-ZnO were encapsulated in the Alg/Plu ink and 3D printed layer by layer with a 3D bioprinter.

**Figure 1.**
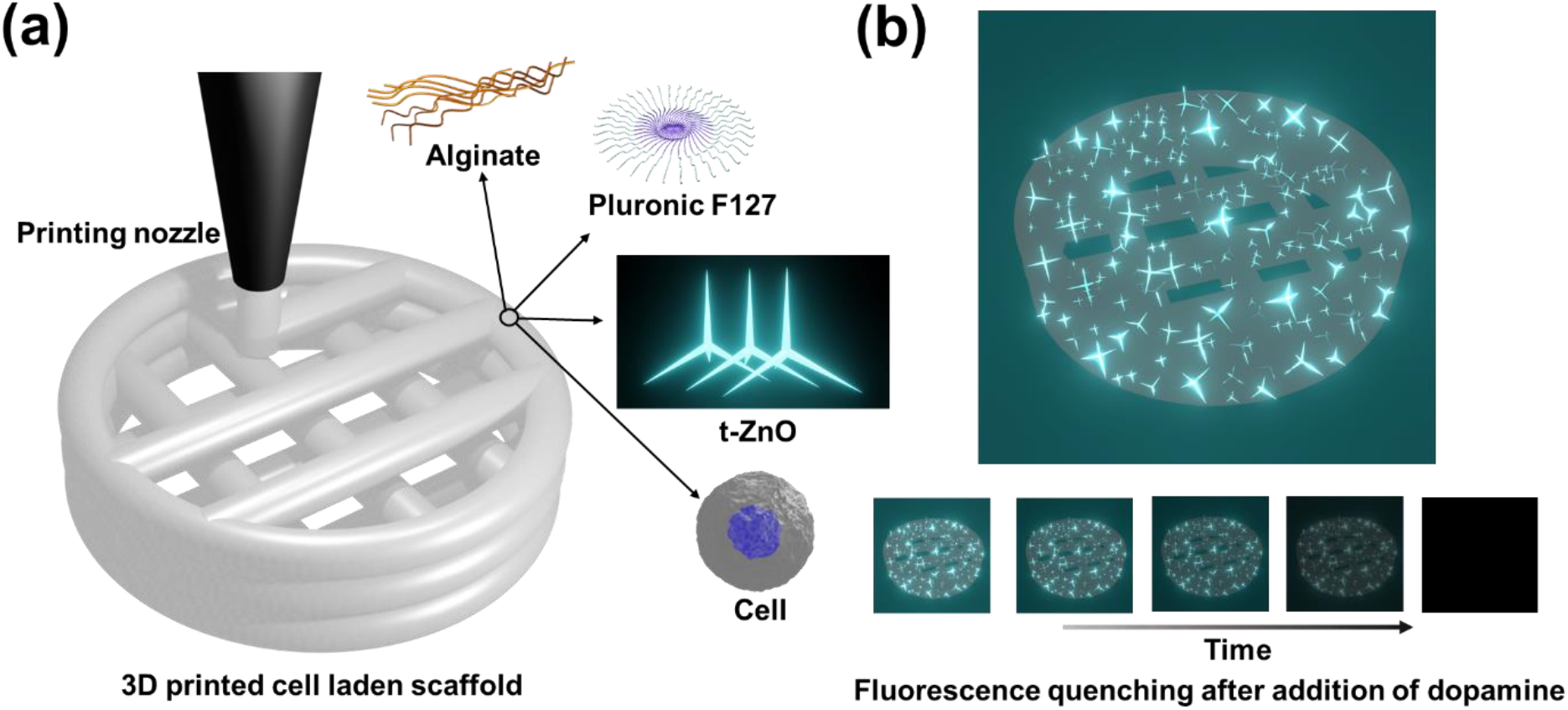
3D printing and autofluorescence of the 3D printed scaffold. (a) Schematic of 3D printing of neural cells in alginate (Alg) and Pluronic F127 (Plu) with t-ZnO additive. (b) top: Autofluorescence of a 3D printed scaffold; bottom: fluorescence quenching of 3D printed t-ZnO with DA over time.

t-ZnO structures used in the experiments had a variety of arm lengths, ranging from 13.2 to 116.0 μm with an average arm length of 48.1 μm (n = 100) (**Figures 2a-b and Figure S1, Supporting Information**). With an excitation at a wavelength of 385 nm, t-ZnO structure emitted fluorescence centered at a wavelength of 536 nm with an intensity positively proportional to the excitation intensity (**Figures 2c-d**). For comparison, with excitation at the same wavelength, ZnO powder (particle size < 5 μm) and glass substrate have no emission (**Figure S2, Supporting Information**), proving the visible emission is an intrinsic property of t-ZnO.

**Figure 2.**
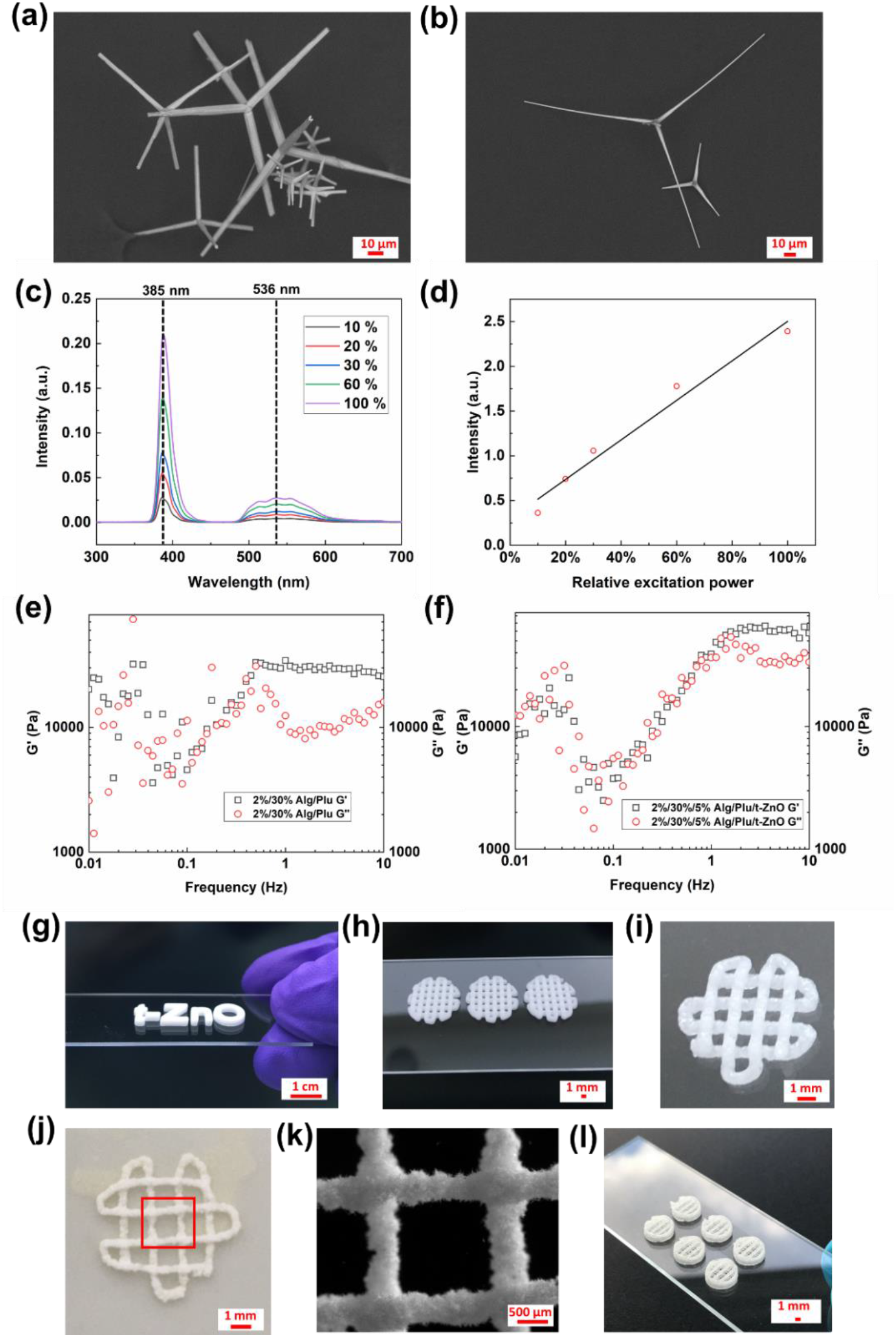
Material and scaffold characterization. (a-b) SEM images of t-ZnO used for 3D printing. (c) Excitation and emission spectroscopy of t-ZnO at different relative excitation powers (10% to 100%). (d) Corresponding plot of emission intensity versus the relative excitation power. Rheological measurements of (e) 2%/30% (w/w) Alg/Plu composite, and (f) 2%/30%/5% (w/w) Alg/Plu/t-ZnO composite versus frequency sweep. (g-i) 3D printed 30%/5% (w/w) Plu/t-ZnO scaffolds with different designs. (j) The 3D printed t-ZnO scaffold after sintering. (k) The image of highlighted area in figure (j). (l) 3D printed 2%/30%/5% (w/w) Alg/Plu/t-ZnO scaffolds on glass slide.

Storage modulus (G’) and loss modulus (G’’) are two important indicators for 3D printability of materials. A material is considered elastic and possible for 3D printing when G’ is higher than G’’ [24]. **Figures S3a-b** (**Supporting Information)** and **Figures 2e-f** demonstrate the evolution of G’ and G’’ with a frequency sweep of 2% (w/w) Alg composite, 30% (w/w) Plu composite, 2%/30% (w/w) Alg/Plu composite, and 2%/30%/5% (w/w) Alg/Plu/t-ZnO composite, respectively. G’’ of 2% (w/w) Alg composite was consistently higher than G’ during the sweeping frequency, which indicates that the 2% (w/w) Alg composite behaved like a fluid rather than a gel (**Figure S3a**). For the 30% (w/w) Plu composite, G’ was higher than G’’ from 0.9 to 3 Hz, indicating its capability to store energy elastically during the frequency range (**Figure S3b**). Interestingly, the 2%/30% (w/w) Alg/Plu composite showed higher G’ and G’’ than 2% (w/w) Alg composite and 30% (w/w) Plu composite over the whole testing frequency, with G’ > G’’ when the frequency was between 0.5 Hz and 10 Hz **(Figure 2e)**. The addition of 5% (w/w) t-ZnO into the 2%/30% (w/w) Alg/Plu composite has decreased the frequency range for G’ > G’’, nevertheless the 2%/30%/5% (w/w) Alg/Plu/t-ZnO composite behaved like gel at high frequency (1-10 Hz) (**Figure 2f**). The result suggests that 2%/30% (w/w) Alg/Plu composite and 2%/30%/5% (w/w) Alg/Plu/t-ZnO composite can be 3D printed at room temperature (RT; 23 °C) with efficient crosslinking.

To create inks for sensor-only scaffolds without any cells, t-ZnO was dispersed in liquidized Plu to maintain a uniform distribution after gelation at RT. The as-prepared 30%/5% (w/w) Plu/t-ZnO ink can be generally used for 3D scaffold printing (**Figures 2g-i**). To obtain pure t-ZnO scaffolds, the printed structures were sintered at 1000 °C (3 h) to remove the organic components (**Figures 2j-k**). This fabrication approach can be applied to create 3D t-ZnO constructs for applications beyond DA sensing^[18]^, e.g., for the fabrication of lightweight framework structures from one-dimensional (1D) and 2D nanomaterials [25-27], as well as for microengineering of functional hydrogels towards tissue engineering [28, 29].

Since Plu is water soluble, for t-ZnO scaffolds that were immersed in aqueous environments and printed with living cells, we added biocompatible Alg to the bio-ink (2%/30%/0.025% (w/w) Alg/Plu/t-ZnO with a cell density of 3 × 10^6^ per mL). The Alg serves as a crosslinkable network after 3D printing to hold the t-ZnO particles structurally stable (**Figure 2l and Video S1, Supporting Information**). t-ZnO distributed homogeneously throughout the 3D printed scaffold with a concentration of 0.025% (w/w) and the printed construct is transparent for fluorescence microscopy, possible for 3D imaging and sensing (**Figure S4, Supporting Information**). The high-throughput 3D printing strategy enable stable and porous t-ZnO scaffold fabrication with potential for applications in various fields.

### 2.2. Sensing Mechanism

In this section, we describe how t-ZnO can be used as an optical sensor for DA. As a catecholamine neural transmitter, DA has both catechol and amine groups that are essential for efficient self-polymerization-based coatings on various surfaces in aqueous media [30]. The polymerization generally occurs in an alkaline pH environment, while an oxidant could also be utilized for inducing the reaction in acidic and neutral media. In our study, t-ZnO can oxidize the catechol moiety of DA to form quinone in the near neural physiological pH, which subsequently leads to the formation of PDA on the surface of the t-ZnO [30, 31]. As an indication of this reaction, a solution of DA in water was found to be stable without any color change, while the mixture of t-ZnO and 1 mM DA became completely dark after two days at RT (**Figure S5, Supporting Information**). Furthermore, 1 mM DA in PBS solution could efficiently quench the t-ZnO fluorescence within 1 hour in a cell culture incubator (**Video S2, Supporting Information**). The deposited amount of PDA was proportional to the DA concentration in the media (**Figures 3a-d**).

**Figure 3.**
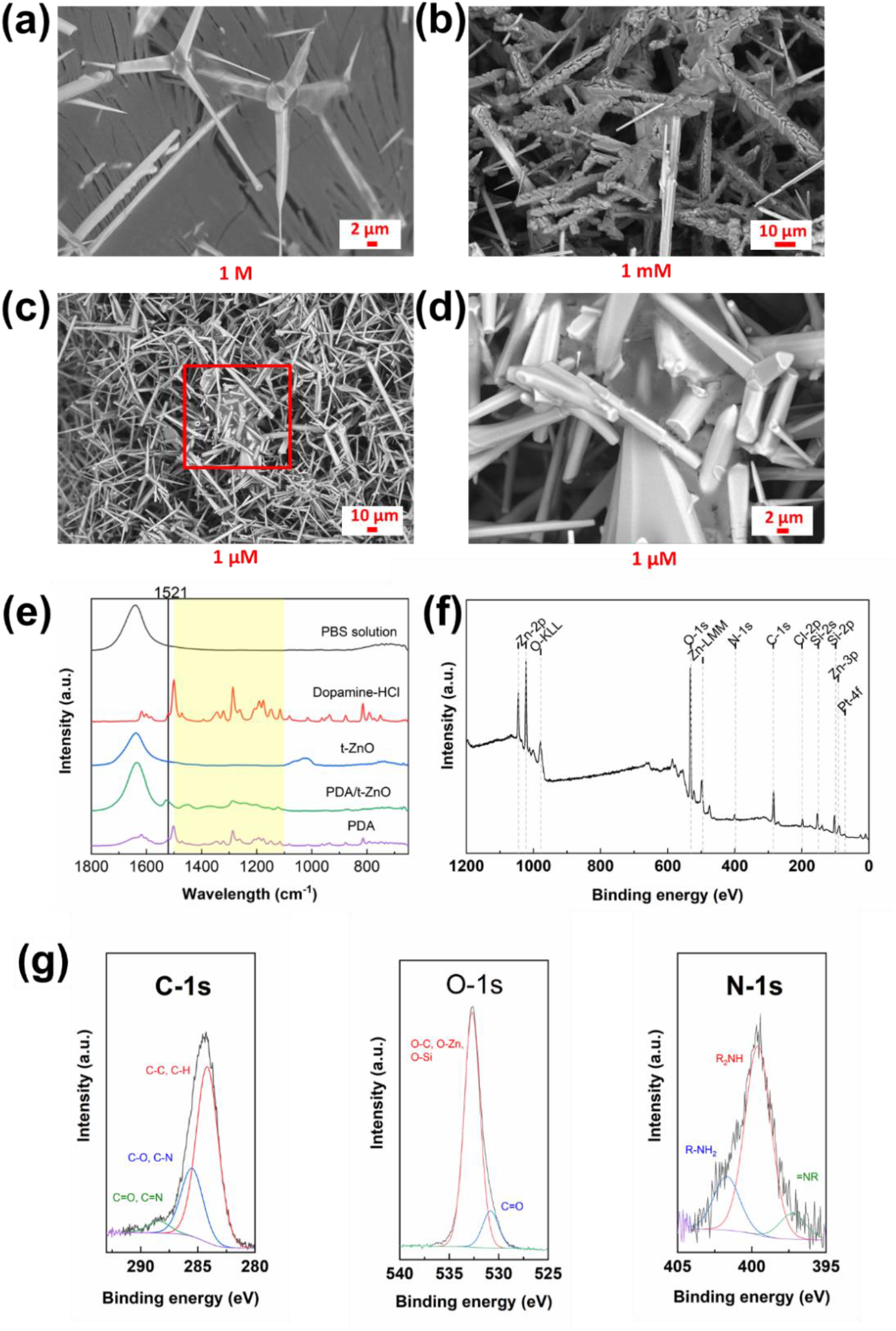
Sensing mechanism investigation. SEM images of polymerized DA (PDA) deposition on the t-ZnO with different DA concentrations of (a) 1 M, (b) 1 mM and (c, d) 1 μM at different magnifications. (e) Fourier transform infrared (FTIR) spectra of phosphate buffered saline (PBS) solution, DA-hydrochloride, t-ZnO, PDA/t-ZnO, and PDA. (f) X-ray photoelectron spectroscopy (XPS) survey scan spectrum of PDA coated t-ZnO shows the presence of O, C, Zn, Si, Pt, N and Cl. (g) High-resolution XPS spectra of the C-1s line, O-1s line and N-1s line.

We carried out Fourier transform infrared (FTIR) and X-ray photoelectron spectroscopy (XPS) to verify the chemical composition of the coatings deposited on the t-ZnO surface. As shown in **Figure 3e**, the DA-HCl spectrum is consistent with the reported results [32]. The peaks at 1521 cm^-1^ are attributed to the aromatic moieties in the structure, while the peaks in the DA-HCl, PDA/t-ZnO and PDA spectra in the yellow highlighted region are ascribed to the C-C, C-N and C-O bonds. The peaks in the spectrum of PDA/t-ZnO in the yellow region are similar to the PDA spectrum, indicating that PDA formed on the t-ZnO surface. As shown in the spectrum of PDA coated t-ZnO (**Figure 3f**), elemental O, C, Zn, Si, Pt, N and Cl were present [33, 34]. All of the peaks belong to the PDA coated t-ZnO sample, except for Si and Pt, which were from sample preparation and testing procedure. To investigate the chemical bonds in the PDA coating layer, high resolution scans of C, O and N were conducted and shown in **Figure 3g**. For comparison, high-resolution scans of Pt, Cl, Zn, Si and overview spectrum of pristine t-ZnO are shown in **Figures S6a-b (Supporting Information)**. The C-1s spectrum was deconvoluted into three peaks, which are corresponding to C-C, C-H bonds (284.2 eV), C-O, C-N bonds (285.5 eV) and C=O, C=N bonds (288.4 eV). These functional groups suggest that DA polymerized on the t-ZnO surface [35], where both catechol and quinone groups were present [36]. The O-1s spectrum could be deconvolved into two peaks. One is attributed to the O-C, O-Zn, O-Si bonds (532.7 eV) and the other to the C=O bonds (530.8 eV). The high-resolution spectrum of the N-1s line was deconvolved into three peaks, which are respectively attributed to =N-R (tertiary/aromatic amine functionalities), R2-NH (secondary amine functionalities) and R-NH2 (primary amine functionalities) at 397.2 eV, 399.6 eV and 401.7 eV [36]. Meanwhile, the N/C molar ratio was measured and calculated to 0.1, which is close to the previously reported stoichiometric value of PDA (N/C= ∼ 0.1) [36]. Therefore, both FTIR and XPS results verify the deposition of PDA on the t-ZnO surface in presence of DA.

Next, we analyze the relationship between the initial DA concentration and autofluorescence quenching. When pristine t-ZnO particles are excited with UV light, they emit fluorescence with wavelengths ranging from 490 to 610 nm, and also dissipates energy non-radiatively (e.g., as heat). This ideal emissive process can be described by

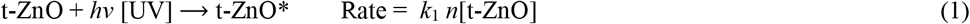

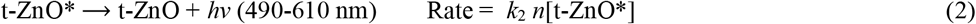

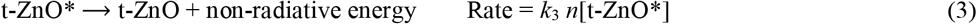

where k_1_, k_2_ and k_3_ are the corresponding reaction rate constants. When the excitation and fluorescence reach steady state, then

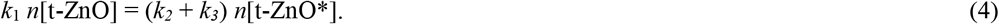

The quantum yield of this process is defined as:

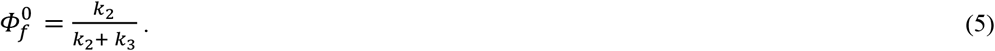

If PDA is adhered onto the t-ZnO surface, the fluorescence is quenched, and the process is

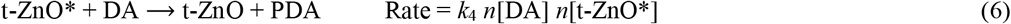

At equilibrium,

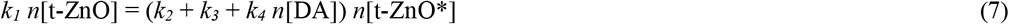

In this case, the quantum yield is given by

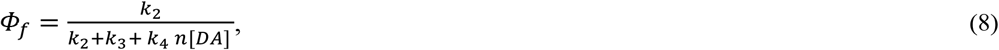

which when combined with equation (Eq) (5) gives

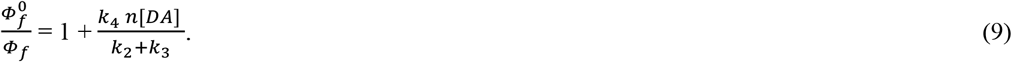

Experimentally, in reference [37] the generation of PDA was found directly proportional to reaction time with a given concentration of DA at the commencement of reaction. This is described by

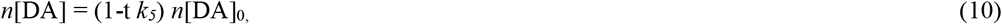

where *k*_*4*_ and *k*_*5*_ are the corresponding reaction rate constant with t as reaction time and *n*[DA]_0_ as a given DA concentration at the beginning of testing. Therefore,

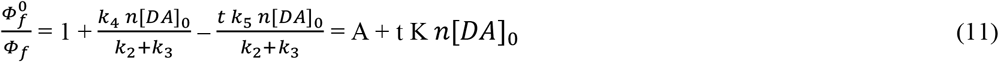

where 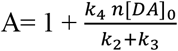 and 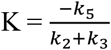 also as a constant. Eq. (11) agrees with the Stern-Volmer relationship [38], which describes the kinetics of quenching. In an experiment, the quantum yield, *Φ*, could be replaced by the fluorescence intensity, *I*. So 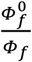 is equivalent to *I*_*0*_*/I*, where *I*_*0*_ and *I* denote respectively the initial and present fluorescence intensity of t-ZnO. According to Equation (11), *I*_*0*_*/I* has a linear relationship with time for a given DA concentration and also has a linear relationship with DA concentration at a fixed time.

### 2.3. Sensor Characterization

The concentration was 1 mM for dopamine, citric acid, glutamine, ascorbic acid, and glucose. The concentrations of KCl and CaCl_2_ were 100 mM and 180 mM, respectively.

To verify the model in the last section, we carried out experiments to investigate the influence of DA on t-ZnO autofluorescence. The time-dependent quenching due to 1 mM dopamine on the emission of t-ZnO is shown in **Figure 4a** and **Video S2 (Supporting Information)**. As the polymerization and deposition of DA on the t-ZnO increased with time, the autofluorescence emission of t-ZnO decreased. With a fixed DA concentration, *I*_*0*_*/I* has a linear correlation with time (R^2^ = 0.9932), as predicted by Eq 11. Similarly, the fluorescence intensity change of t-ZnO as a function of different DA concentrations at a given time (30 min) is shown in **Figure 4b**. A linear correlation between *I*_*0*_*/I* and DA concentration (5 μM – 1000 μM) was observed (y = 1.12 + 6.53 × 10^−4^ x; R^2^ = 0.9938), with a detection limit of 0.137 μM (S/N = 3). Previous ZnO optical DA sensors had a detection limit of 0.791 μM and a reaction time of 6 hours [19, 20]. Thus, the t-ZnO had a 5.8x lower detection limit and was about 12x faster. The higher sensitivity of t-ZnO may be due to its 3D high aspect ratio structure, allowing more active sites for DA reaction.

**Figure 4.**
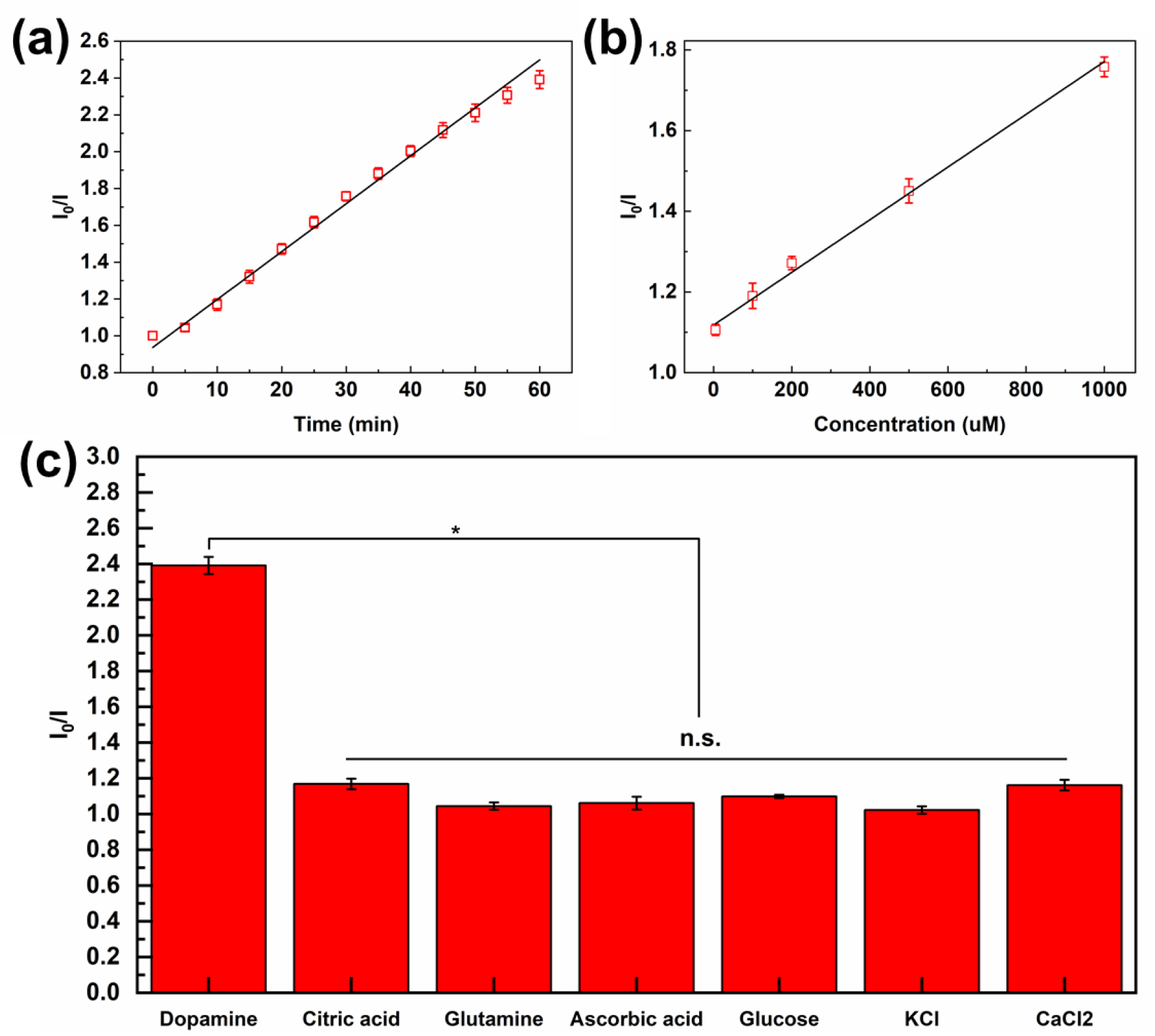
Experimental evaluation of t-ZnO based DA sensor. (a) Time-dependent quenching of the emission of t-ZnO due to DA plotted as *I*_*0*_*/I* versus time. (b) t-ZnO autofluorescence intensity (*I*_*0*_*/I*) as a function of DA concentration after 30 mins. (c) t-ZnO autofluorescence intensity after 60 mins in solutions of common chemicals found in the brain.

We further verified the specificity of DA sensing by exposing the t-ZnO to common neurochemicals and ions present in the brain for 60 min, including 1mM of DA, 1 mM citric acid, 1 mM glutamine, 1 mM ascorbic acid, 1 mM glucose, 100 mM K^+^ and 180 mM Ca^2+^. As shown in **Figure 4c**, *I*_*0*_*/I* value of DA is more than 100% higher than the other interfering molecules and ions, showing high selectivity of DA over other interferences.

### 2.4. Characterization of 3D Printed Neural Tissue with t-ZnO Additive

3D printed Alg based neural tissue is an improved resemble of *in vivo* counterpart [39]. Scaffolds tend to disintegrate during culture due to chelating agents [24, 40], but the reinforcement of the printed scaffold in a crosslinking solution can maintain the mechanical strength and structural integrity after Plu is dissolved. **Figures 5a-c** show fluorescence microscope and scanning electron micrographs of the cell morphology after 3D printing. The results confirmed that the 3D printing SH-SY5Y cells with t-ZnO and reinforcement with CaCl_2_ solution during culture did not negatively influence the cell viability and 3D neural network forming (**Figures 5a-c)**. After 7 days of culture, the 3D printed scaffold continued to maintain its structural integrity with homogenously distributed living cells and t-ZnO in the construct (**Figure 5a**). SEM images and immunostaining images revealed extensive interconnected 3D neural network projections with typical dopaminergic neuron morphology formed across the 3D structure from day 1 to day 7, without any apparent negative effects from the t-ZnO (**Figures 5b-c and Figures S7-9, Supporting Information**). DA released from a synapse could polymerize on the t-ZnO surface, which subsequently quenches the autofluorescence of t-ZnO to indicate the dopaminergic tissue activity as discussed in the previous section. Immunostaining also indicated that neuronal marker β-tubulin III and the key cytoskeletal component F-actin have increased significantly on day 7 compared to day 1 (**Figure 5c and Figure S9, Supporting Information)**. Co-expression of β-tubulin III and F-actin shows that microtubule and F-actin serve important role in forming neuronal networks and powering growth cone motility [41]. The extensive expression of F-actin in the neural projects indicates that the ink with t-ZnO did not influence growth cone moving and dendritic spine dynamics, suitable for artificial neural tissue engineering [41, 42].

**Figure 5.**
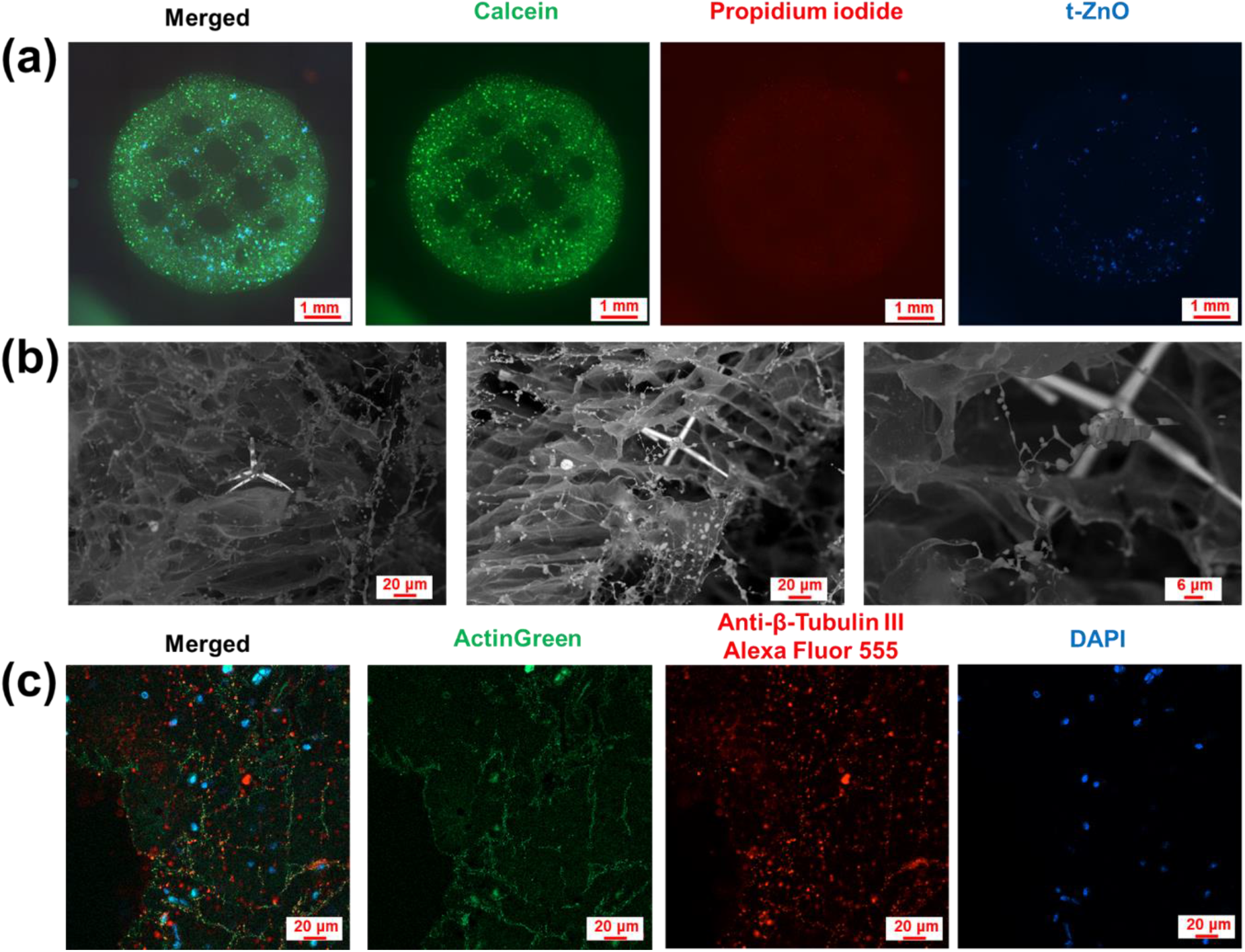
Cytocompatibility and morphology of 3D printed SH-SY5Y cells with embedded t-ZnO 7 days after printing. (a) Fluorescence microscope images of 3D printed SH-SY5Y cells with t-ZnO. Live (Calcein; green), dead [propidium iodide (PI); red] and t-ZnO (autofluorescence; blue). (b) Representative SEM images of 3D printed cells with t-ZnO at different locations and magnifications. (c) Confocal fluorescence microscope images of fixed 3D printed SH-SY5Y cells with t-ZnO. F-actin staining in green, immunostaining of neuronal marker β-tubulin III in red, and nuclei staining in blue.

### 2.5. Sensor Characterization in Other Environments

The t-ZnO sensor was used for DA sensing in the 3D printed dopaminergic tissue. However, different physiological environments, especially those with metallic ions, can have an impact on the polymerization of DA on the t-ZnO surface) [43]. To compare the sensing results in PBS, we used a differentiation medium (DM) as an alternative environment for the sensor. A dilution ratio of 10 with PBS gives significantly high recovery (107%) compared to pure DM (66%), almost equivalent to the result from pure PBS platform (106%) (**Table S1, Supporting Information**). The same diluted DM also gave similar recoveries to other spiked DA concentrations (600 μM, 200 μM, and 50 μM), showing that the efficiency of t-ZnO sensor can be maintained in a variety of aqueous environments (**Table S2, Supporting Information**).

Lastly, we used t-ZnO as an *in situ* sensor to evaluate the dopaminergic cell activity in the 3D printed neural tissue. Not only can t-ZnO serve as a qualitative indicator of DA release, it can also be used to quantify the dopamine release if the intensity is calibrated. t-ZnO embedded in the 3D printed neural tissue fluoresced when excited by a wavelength of 385 nm (**Figures 6a-b**). Three regions of interest (ROIs) were selected at different depths of a t-ZnO structure and its autofluorescence was measured over 30 minutes. In **Figure 6a**, *I*_*0*_*/I* values of the three ROIs are 1.08, 1.12 and 1.14, indicating ROI 2 and ROI 3 have more dopaminergic activities then ROI 1. Similarly, *I*_*0*_*/I* values of three ROIs in **Figure 6b** are 1.03, 1.02 and 1.00, indicating much lower dopaminergic activity compared to the ROIs in **Figure 6a**. When *I*_*0*_*/I* value is larger than 1.12 (y interception of fitted equation *y* = 1.12 + 6.53 × 10^−4^*x* from **Figure 4b**), the released DA can be estimated with the equation. For example, ROI 3 in **Figure 6a** indicates the regional DA concentration was about 46 μM (Also taking the recovery of DA in pure DM from **Table S1** into consideration). For *I*_*0*_*/I* < 1.12, a longer testing time could be used for quantitative analysis. Accordingly, the DA time dependence can be measured in 3D spatially by tracing the *I*_*0*_*/I* value of the embedded t-ZnO particles (**Figures S11-S12, Supporting Information**). Using optical imaging to detect DA enables the 3D sensor to have a micron-scale spatial resolution and time resolution of seconds, but the spatial resolution is limited by the density of the t-ZnO particles and the imaging optics. In our experiments, in a plane, t-ZnO occupied about 11.3 % (± 0.4 %) of the area of the printed structure, which means t-ZnO particles were distributed about with an estimated average separation of about 150 μm. To have higher coverage in the 3D scaffold, higher t-ZnO content, and a higher spatial resolution can be achieved with a higher magnification objective lens. It is also possible to position a t-ZnO structure at a specific location after tissue printing or on tissue to monitor the local DA concentration.

**Figure 6.**
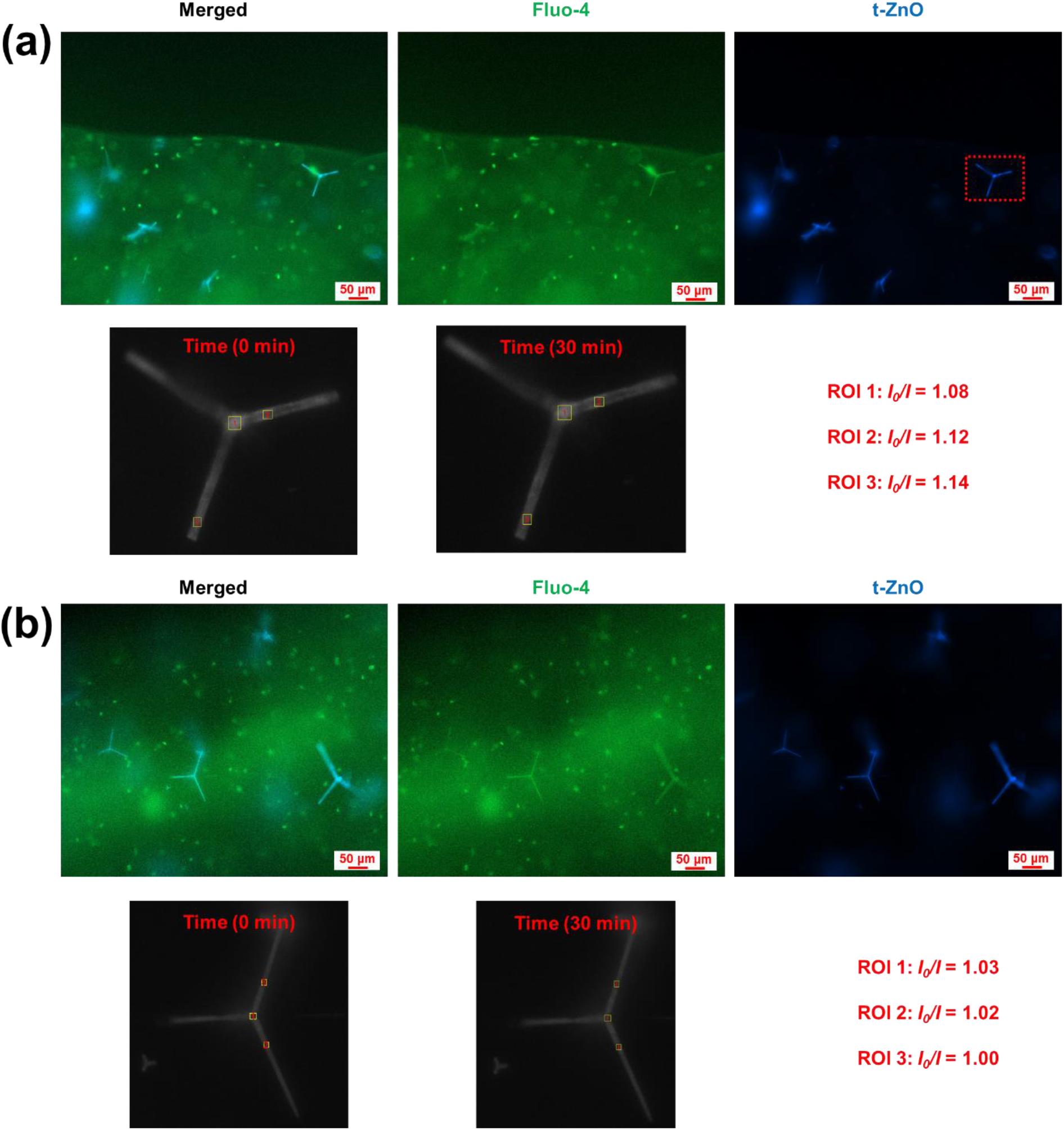
DA sensing in the 3D printed neural tissue 7 days after printing. Fluorescence microscope images of 3D printed SH-SY5Y cells with t-ZnO for 30 min (a) at the surface of structure (depth: 0 μm) and (b) in the middle of the structure (depth: 100 μm). 3 region of interests (ROIs) were selected for each t-ZnO. Cell (Fluo-4; green) and t-ZnO (autofluorescence; blue).

## 3. Conclusion

In summary, we have presented a novel method for 3D neuron tissue bioprinting with SH-SY5Y cells as a dopaminergic model and t-ZnO as an *in situ* DA sensor. t-ZnO is cytocompatible and its autofluorescence is quenched by the polymerization of DA, in which t-ZnO oxidized the catechol moiety of DA at physiological pH and catalyzed the subsequent polymerization process on the surface of t-ZnO. No surface functionalization or modification is needed. The dynamics of the quenching were found to agree with Stern-Volmer relationship theoretically and experimentally. The t-ZnO based sensor has high selectivity of DA over other interfering molecules and a linear detection range from 5 to 1000 μM with a detection limit of 0.137 μM. Moreover, the developed bio-ink could also be used to 3D print pure t-ZnO scaffold by high temperature sintering after printing. t-ZnO could be used to quantitatively monitor the dopamine release with cellular resolution. An ultimate goal of tissue engineering is to fabricate clinically relevant tissues for translational medicine screening, human tissue/disease development studies, or replacement of diseased human tissue or organ. Thus far, effective *in situ* and contactless methods of monitoring the cells in the engineered tissue or organ are lacking. We have shown that neurochemical sensors can be integrated with tissues in 3D, paving the way for the development of smart tissues with optical monitoring.

## 4. Experimental Section

### Materials

Sodium alginate from brown algae (Alg, average Mn: 80,000-120,000), Pluronic F-127 (average Mn: 12,600), dopamine hydrochloride (DA-HCl), zinc oxide powder (particle size < 5 μm), citric acid monohydrate, L-ascorbic acid, calcium chloride (CaCl_2_) dihydrate, retinoic acid (RA), Calcein AM and propidium iodide (PI) were obtained from Merck (Germany) and used as received. 3D printing syringe barrels and needles were purchased from Nordson EFD (United States). Gibco*™*L-glutamin (200 mM), Gibco*™*penicillin-streptomycin (10.000 U/ml), Cytiva HyClone*™*newborn bovine calf serum (heat inactivated) was purchased from Fisher Scientific (Germany). D-(+)-Glucose anhydrous was purchased from Molekula group (France). Dulbecco’s Modified Eagle’s Medium (DMEM) was obtained from Corning (USA). Glass slide and cover slip were obtained from VWR (Germany). Highly pure and single crystalline t-ZnO was produced with a previously reported flame transport synthesis method [44].

### Ink preparation

2% (w/w) Alg solution was prepared by adding 200 mg Alg in 10.000 g water with constant stirring and heating (80 °C) for 3 h. 30% (w/w) Plu solution was prepared by mixing 3.000 g Plu in 10.000 g water with mechanical mixing in the fridge (4 °C) for 10 h. 3.000 g Plu was dissolved in the prepared 2% (w/w) Alg solution at 4 °C for 10 h. 500 mg t-ZnO was dispersed in the 2%/30% (w/w) Alg/Plu mixture with constant mechanical stirring. The obtained 2%/30%/5% (w/w) Alg/Plu/t-ZnO mixture was transferred to a 3 mL printing plastic barrel, with subsequent air bubble removal by centrifugation (Eppendorf, Germany). 30%/5% (w/w) Plu/t-ZnO ink was prepared by the same process.

### Rheological measurements

A rotational rheometer (HAAKE™MARS™, Germany) was employed for the rheology tests with a P35/Ti measuring geometry (35 mm in diameter) fitted and an air-cooled Peltier module for temperature control. A working gap distance between the test geometry and plate was 500 µm, and samples were placed on the rheometer plate before moving the test geometry to the working distance with excess samples removed afterwards. Oscillation sweep was performed from 0.01 Hz to 10 Hz at RT to determine the storage modulus (G′) and loss modulus (G′′).

### 3D printing of t-ZnO

3D printed objects were designed in BioCAD (RegenHU, Switzerland) with the generated G-code loaded into the 3D Discovery printer (RegenHU, Switzerland) for printing. 3D t-ZnO scaffolds were printed layer-by-layer with G21 needle (Nordson, USA) at RT. Scaffolds printed by 30%/5% (w/w) Plu/t-ZnO ink were sintered in a furnace (Eurotherm, Germany) at 1000 °C for 3 h to combust the organic component and obtain the pure t-ZnO scaffold. Scaffolds printed by 2%/30%/5% (w/w) Alg/Plu/t-ZnO were crosslinked with the Alg component in 2% (w/v) CaCl_2_ solution for 10 min.

### Scanning electron microscopy

Morphology of t-ZnO was analyzed with a Hitachi TM4000Plus scanning electron microscopy (SEM) (Hitachi, Japan), after depositing of 5 nm platinum layer on the structure with an MC1000 ion sputter coater (Hitachi, Japan). For the 3D printed scaffolds with cells, samples were fixed with 4% paraformaldehyde PBS solution (PFA, ThermoFisher Scientific) supplemented with 5 mM CaCl_2_ for 30 min, rinsed 3 times in PBS solution, and then observed on an ultra coolstage at -30 °C (Deben, UK) with Hitachi TM4000Plus SEM.

### Photoluminescence (PL) spectroscopy

Photoluminescence (PL) spectroscopy of t-ZnO was measured with a compact Thorlabs CCS200 spectrometer (Thorlabs, USA). Samples were dispersed in the water and dried on a glass slide before testing.

### Fourier transform infrared spectra (FTIR)

FTIR were tested and recorded on a Bruker INVENIO-R Spectrometer equipped with a MIRacle Ge-ATR. Dopamine-HCl was tested as received without further treatment, and other samples were processed in phosphate buffered saline (PBS). For each sample, 20 scans were collected with a resolution of 4 cm^-1^ and reference of N_2_.

### X-ray photoelectron spectroscopy (XPS)

XPS (Omicron Nano-Technology GmbH, Al-anode, 240W) was used to characterize the PDA coated t-ZnO. To exclude contamination from other elements, 1 mM dopamine water solution was mixed with t-ZnO for two days and dried afterwards in the fume hood. The obtained powder was pressed onto a Si wafer with native oxide layer and Pt contacts. The samples were pressed to avoid the usage of any kind of glue, which may influence XPS measurements. The Pt contacts were used for subsequent charge calibration. The charging of sample was corrected by calibrating the Pt4f 7/2 line (**Figure S5b, Supporting Information**) to 71.2 eV. CasaXPS (version 2.3.16) was used for the analysis of XPS spectra.

### Sensing characterization

2 mg t-ZnO per well in a 96-well plate was added with 100 μL dopamine PBS solution (concentrations range from 5 μM to 1 mM). For dopamine sensing measurement, an inverted microscope (Zeiss Axio Observer, Germany) with incubation system (37°C and humidified 5% CO_2_; XLmulti S2 DARK, Germany) was used for image taking with excitation light at 385 nm and ImageJ/Fiji software was used for fluorescence intensity extraction [45]^]^. The selectivity test was implemented with the same procedures (1 h incubation time), while 1mM citric acid, 1 mM glutamine, 1mM ascorbic acid, 1mM glucose, 2% (w/w) CaCl_2_, and 100 mM KCl were added as the interfering agents.

### Human SH-SY5Y neuroblastoma cells culture and differentiation

Human SH-SY5Y neuroblastoma cells (Elabscience, China) were grown in cell culture medium within an incubator (37 °C and humidified 5% CO_2_; Binder, Germany). The cell culture medium was prepared with penicillin (20 units/mL), streptomycin (20 ug/mL), 10% (v/v) heat inactivated fetal calf serum and DMEM. A cell density of 1×10^4^ cells/cm^2^ was used for cell subculture. The culture medium supplemented with 10 μM RA was used as differentiation medium (DM) to induce SH-SY5Y cells towards dopaminergic neuron lineage differentiation, which has been widely applied as neurodegenerative disease model [46]. RA can inhibit cell growth and DNA synthesis by arresting cells in G1 phase, so differentiated cells have lower confluency than undifferentiated cells with the same initial cell density [47]. The culture medium was changed every two days.

### 3D printing of SH-SY5Y cells with t-ZnO

To eliminate interference of auto-fluorescence from t-ZnO, lower concentration of t-ZnO was used for 3D printing with SH-SY5Y cells. 1 mL 2%/30%/0.025% (w/w) Alg/Plu/t-ZnO ink was prepared as described in previous sections and 3 × 10^6^ SH-SY5Y cells were added at 4 °C, following with 1 min mechanical mixing at 1000 rpm. The obtained ink was transferred into a 3 mL printing barrel and subsequently centrifuged at a speed of 500 xg for 2 min (4 °C) to remove air bubbles. The ink was incubated in 37 °C water bath for 5 min before being 3D printed. 3D bioprinting was performed with a G21 needle and 3D Discovery printer (RegenHU, Switzerland). The printed scaffolds were ionically crosslinked in copious 2% (w/w) CaCl_2_ for 1 min and rinsed with fresh culture medium before being transferred into incubator for long term culture. Crosslinked Alg scaffold became gradually loosed or even dissolved in the culture medium due to monovalent cation replacement of divalent calcium cation. To maintain the structural integrity, scaffold was reinforced in 1.5% (w/w) CaCl_2_ for 1 min the day after being printed and before viability testing.

### Cytocompatibility staining assay

Calcein AM and PI were used as cytocompatibility staining assay at a final staining concentration of 4 μM and 2 μM, respectively, to evaluate cell survival and distribution in the 3D cell culture. Samples were washed in fresh culture media after 30 min staining in the incubator before imaging with an Axio Observer microscope (Zeiss, Germany).

### Immunostaining assay

Samples were fixed with 4% PFA PBS solution for 30 min at RT, rinsed 3 times in PBS solution, and followed by blocking and permeabilization with 10% (v/v) horse serum (HS, MP Biomedicals) in PBS supplemented with 0.3% (v/v) Triton X-100 (Calbiochem) at RT. After blocking and permeabilization, samples were washed 3 times in PBS and subsequently incubated with primary mouse anti-β-tubulin III antibody (1:100; Merck) in 10% (v/v) HS PBS solution overnight at 4 °C. On the second day, samples were rinsed with PBS 3 times and then incubated with Alexa Fluor 555 conjugated secondary goat anti-mouse IgG antibody (1:100; Merck) for 2 hours at RT. F-actin and nuclei were stained with 2 drops/mL AlexaFluor*™*488 phalloidin (ThermoFisher Scientific) and 2 µg/mL 4′,6-diamidino-2-phenylindole (DAPI; Merck) for 1 hour at RT, respectively. After staining, samples were washed 3 times in PBS and then mounted onto a glass coverslip (Fisher Scientific) with ProLong*™*glass antifade mountant (ThermoFisher Scientific). Samples were imaged with a confocal microscope (Zeiss LSM 900) and analyzed with Zeiss Zen software. To prevent crosslinked Alg dissolution, all the PBS solutions used were added with 5 mM CaCl_2_.

### Data and statistical analyses

OriginPro 2019 software was employed for data statistical analyses and graphing. All experiments were performed in triplicate and the obtained quantitative data were presented as mean ± standard deviations (SD). A one-way ANOVA analysis with Bonferroni test was used for statistical difference examination and statistical significance was indicated when *P* is smaller than 0.05.

## Acknowledgements

This work is financially supported by Max Planck Society. RA and FS acknowledge funding by the German Research Foundation (Deutsche Forschungsgemeinschaft, DFG) through the RTG 2154. JL would like to thank Dr. Yiding Lin, Dr. Youngho Jung, Mr. Fu-Der Chen, Dr. Andrei Stalmashonak, Mr. Frank Weiß, Ms. Heike Menge and Dr. Daniel Meyer for helpful discussion and technical support.

