## Supplemental document for "3D Printed Neural Tissues with *in situ* Optical Dopamine Sensors"

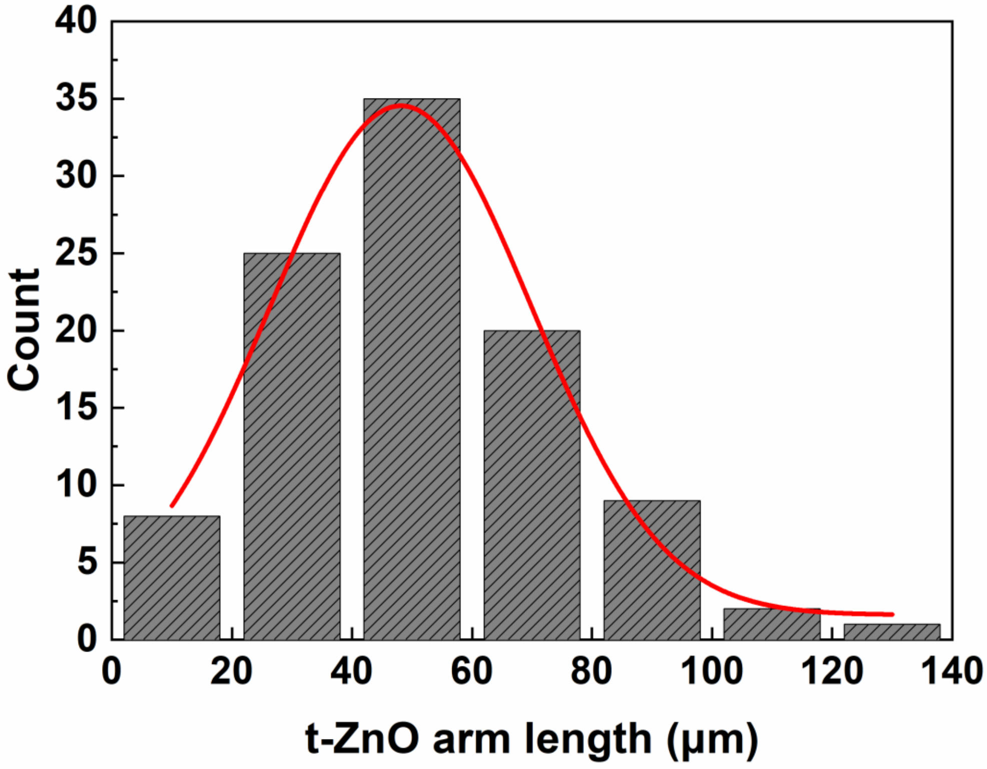


**Figure S1.** Histogram of the t-ZnO arm lengths distribution as extracted from 6 SEM images.

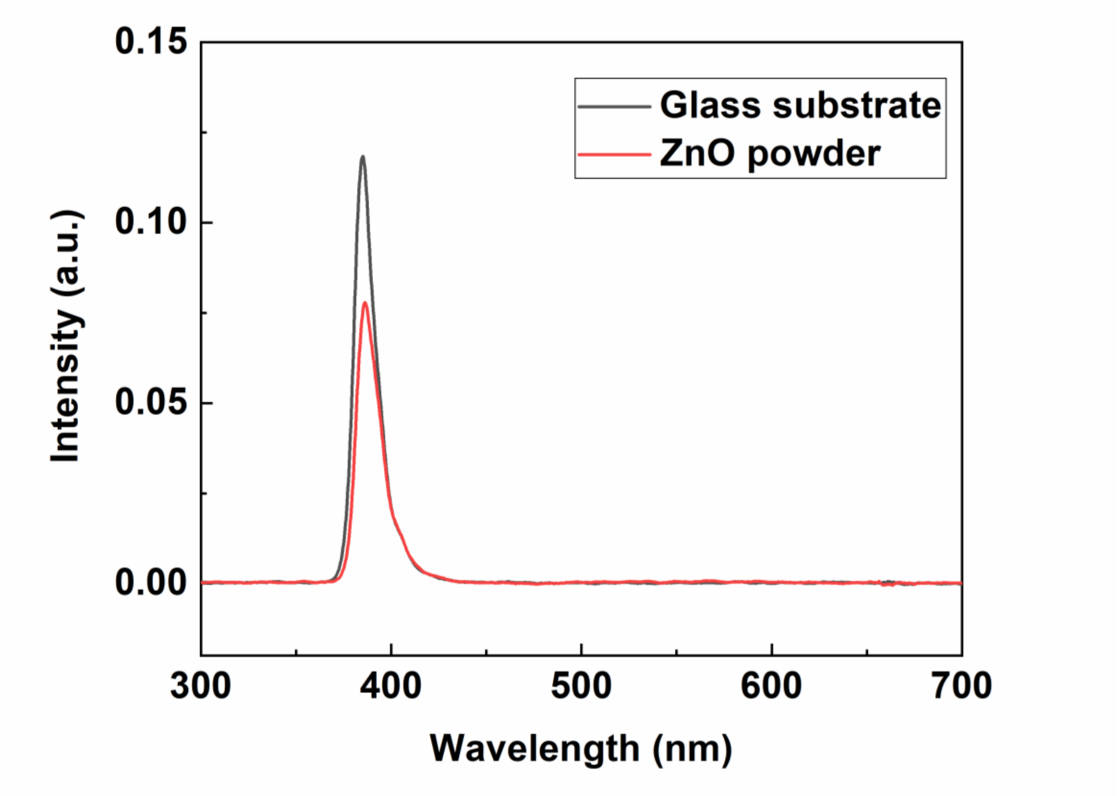


**Figure S2.** Excitation and emission spectra of glass substrate and ZnO powder at 25% excitation intensity.

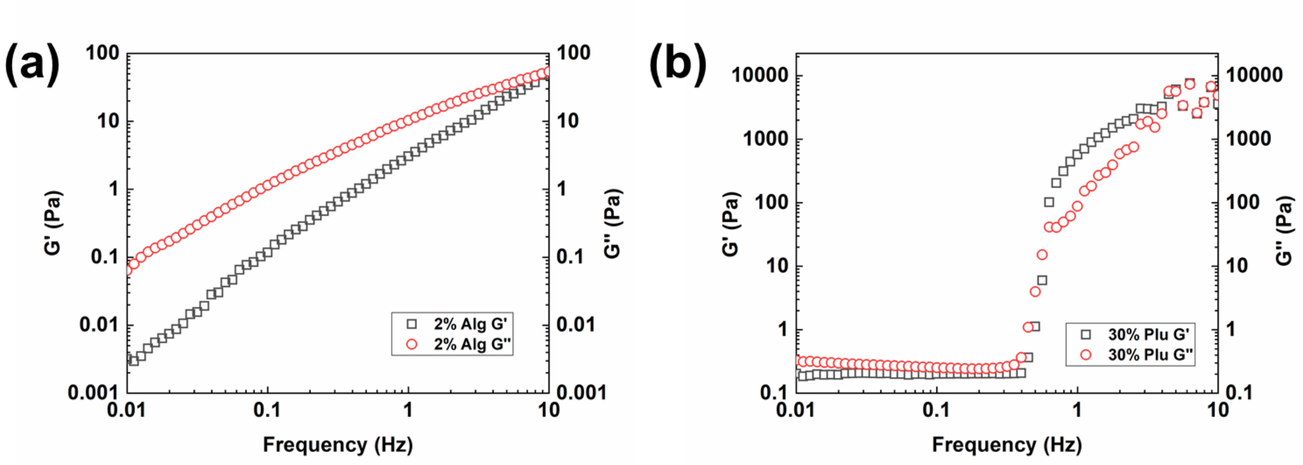


**Figure S3.** Rheological measurements of (a) 2% (w/w) Alg composite, b) 30% (w/w) Plu composite versus frequency sweep.

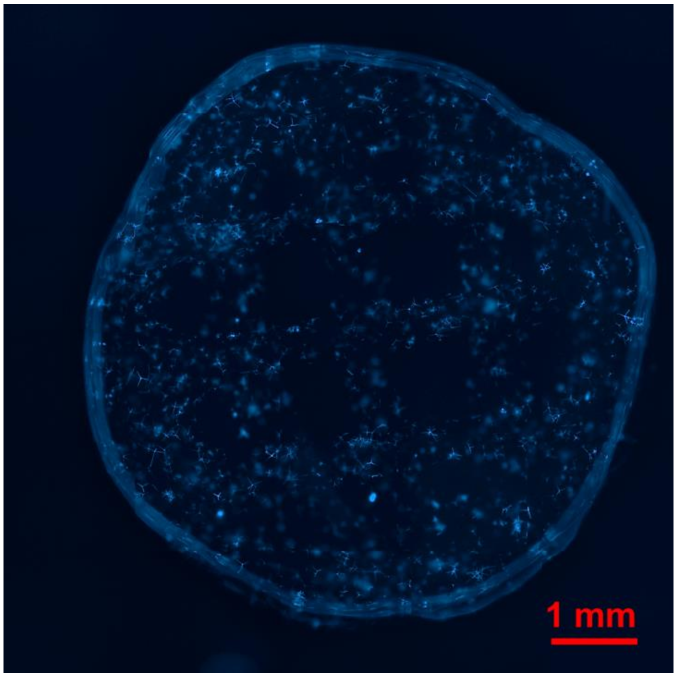


**Figure S4.** Fluorescence microscopy image of 3D printed 2%/30%/0.25% (w/w) Alg/Plu/t-ZnO scaffold.

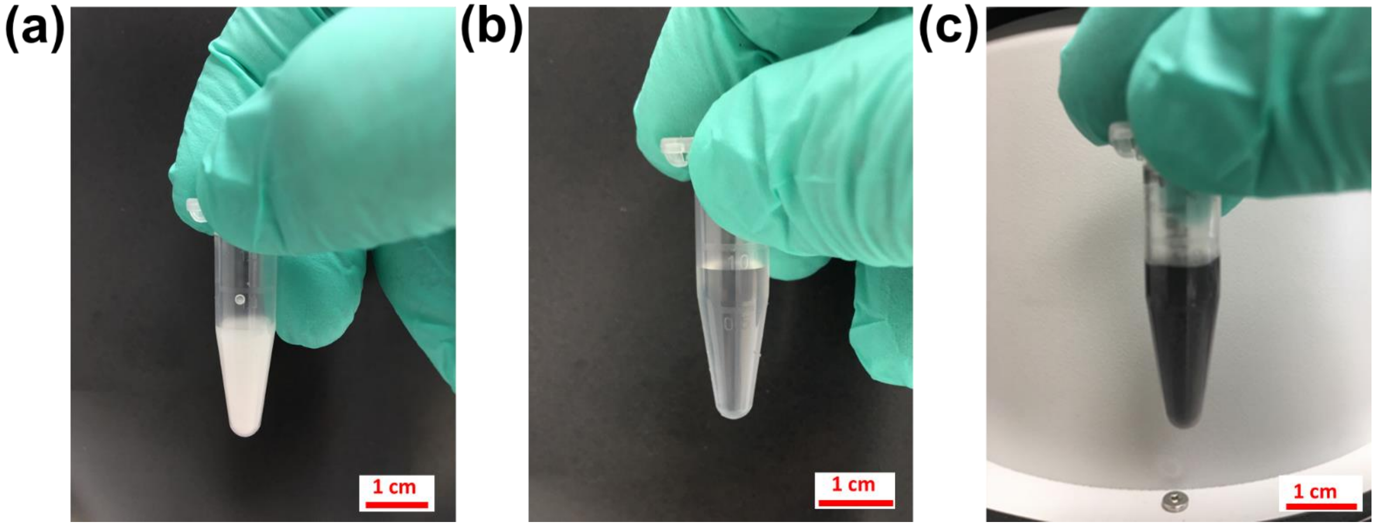


**Figure S5.** Dopamine polymerization with t-ZnO. Images of (a) t-ZnO dispersed in water, (b) 1 mM dopamine water solution and (c) t-ZnO dispersed in 1 mM dopamine solution after incubation for 2 days at room temperature, correspondingly.

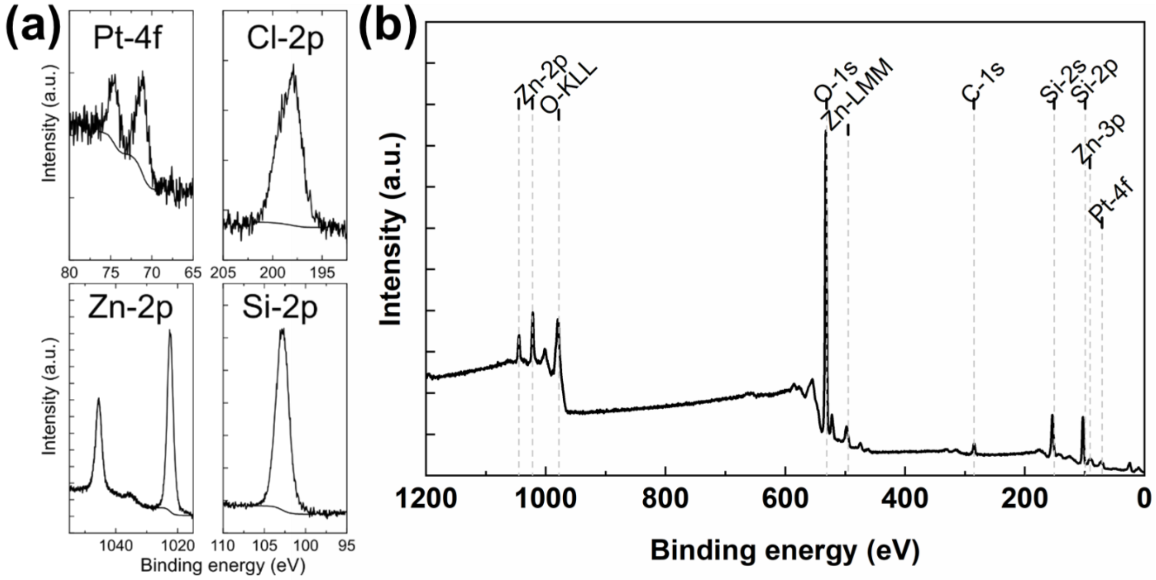


**Figure S6.** (a) High resolution XPS spectra of Pt-4f, Cl-2p, Zn-2p and Si-2p lines from PDA coated t-ZnO measurement. The Pt-4f line was used for calibration. (b) XPS survey scan spectrum of pristine t-ZnO.

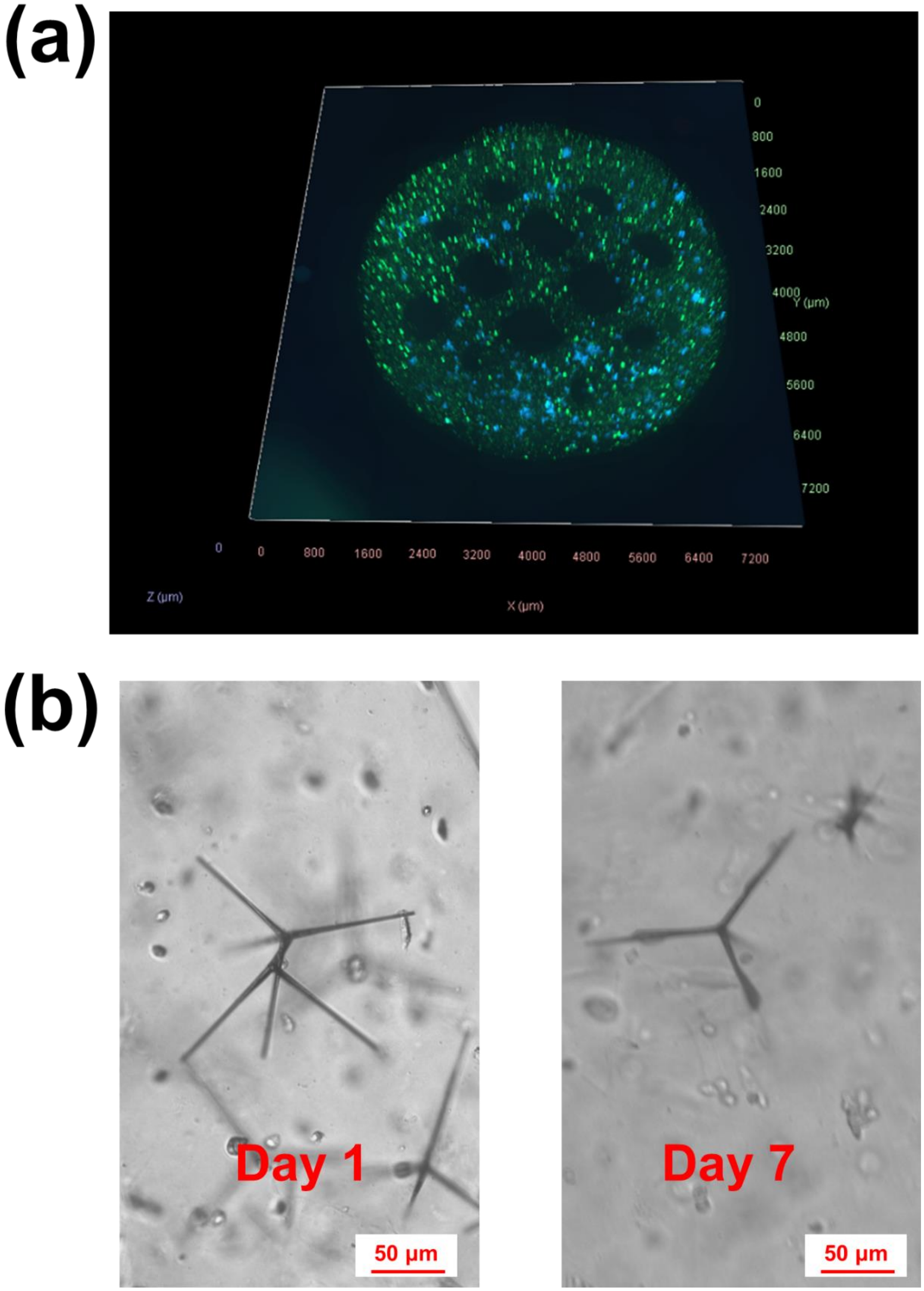


**Figure S7.** (a) 3D fluorescence microscope image of printed SH-SY5Y cells with t-ZnO 7 days after printing. Live (Calcein AM; green), dead (propidium iodide; red) and t-ZnO (autofluorescence; blue). (b) Bright field images of 3D printed SH-SY5Y cells 1 day and 7 days after printing, respectively.

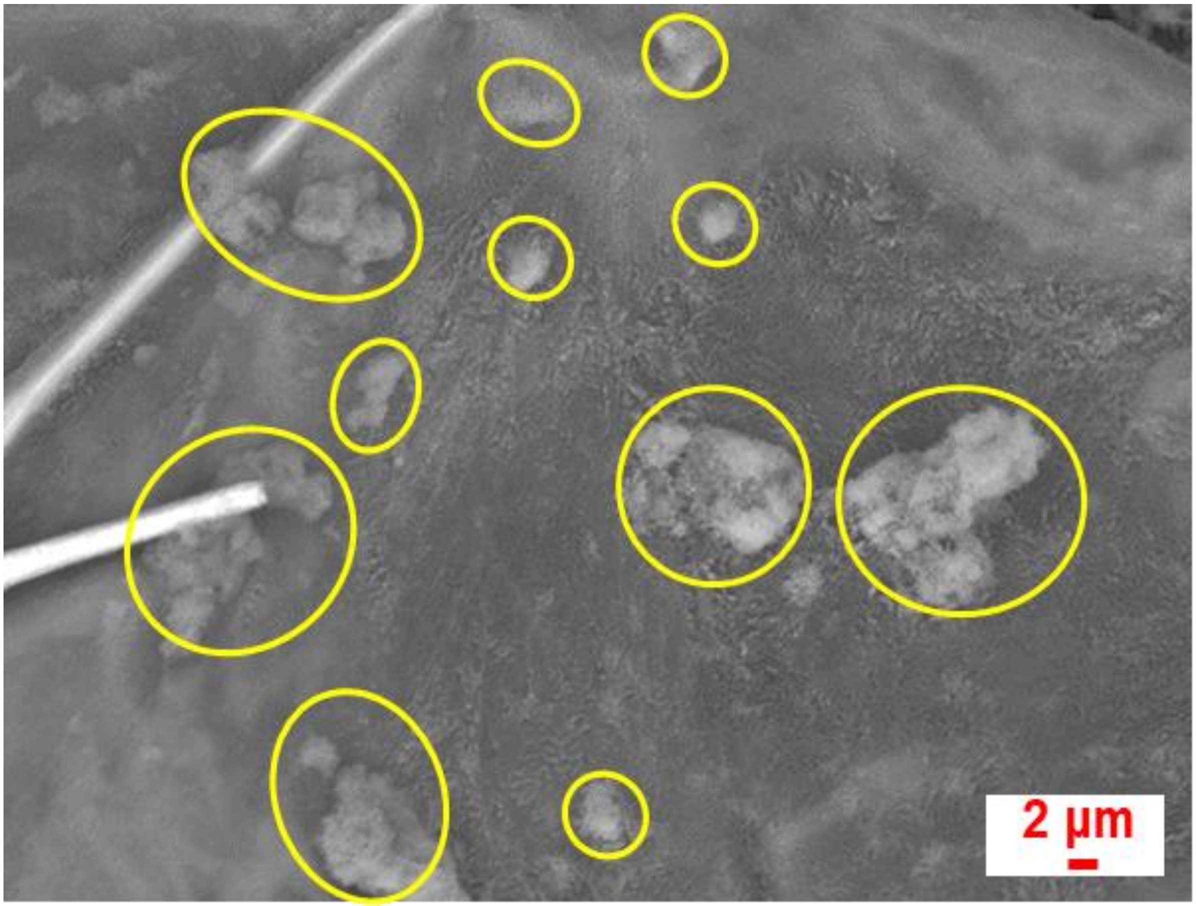


**Figure S8.** Representative SEM image of 3D printed cells with t-ZnO 1 day after printing. Cells are identified with yellow circles.

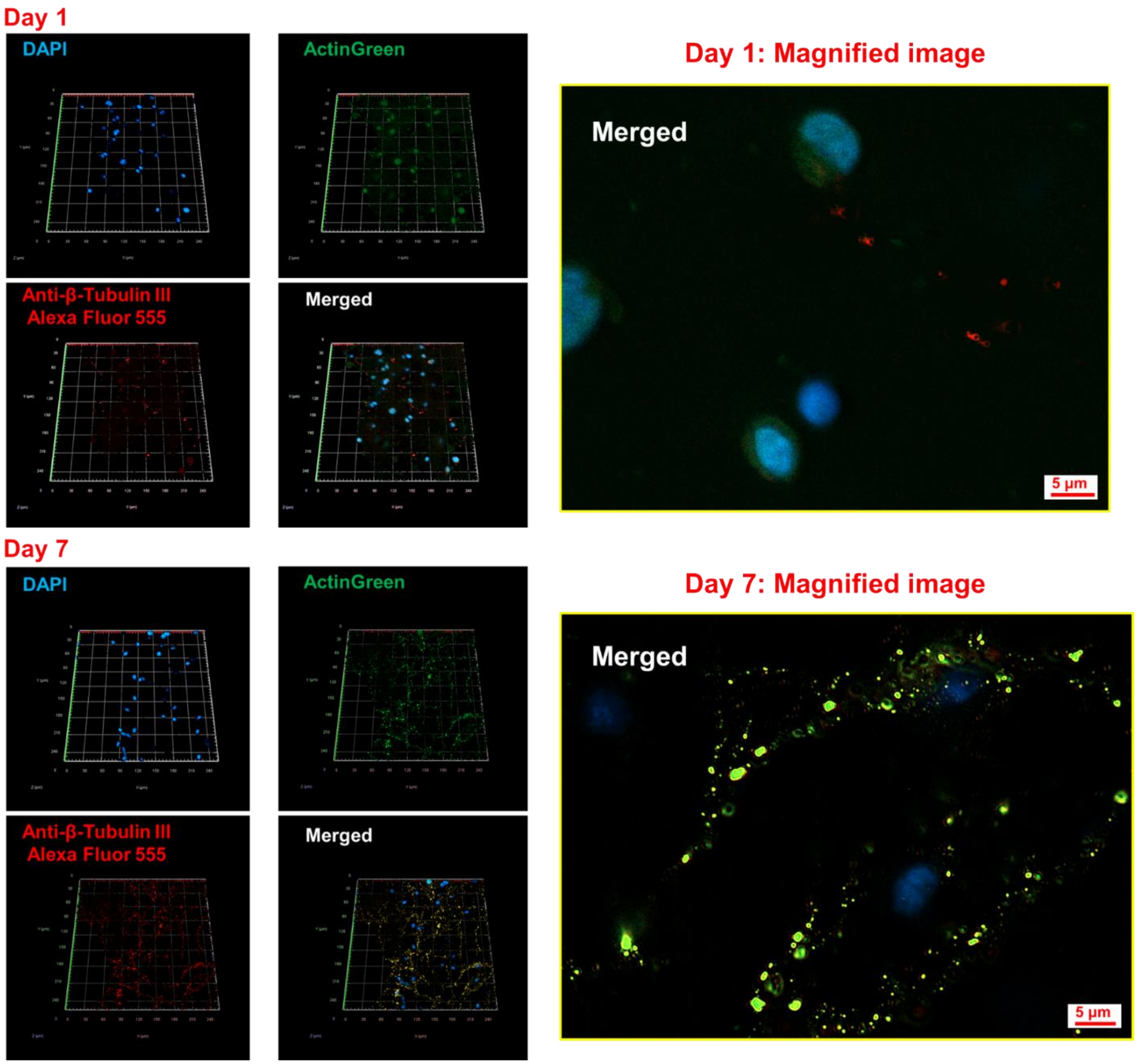


**Figure S9.** Morphology of 3D printed SH-SY5Y cells with t-ZnO after 1 day and 7 days culture respectively. Nuclei (DAPI; blue), F-actin (ActinGreen; green) and β-tubulin III (Alexa Fluor; red).

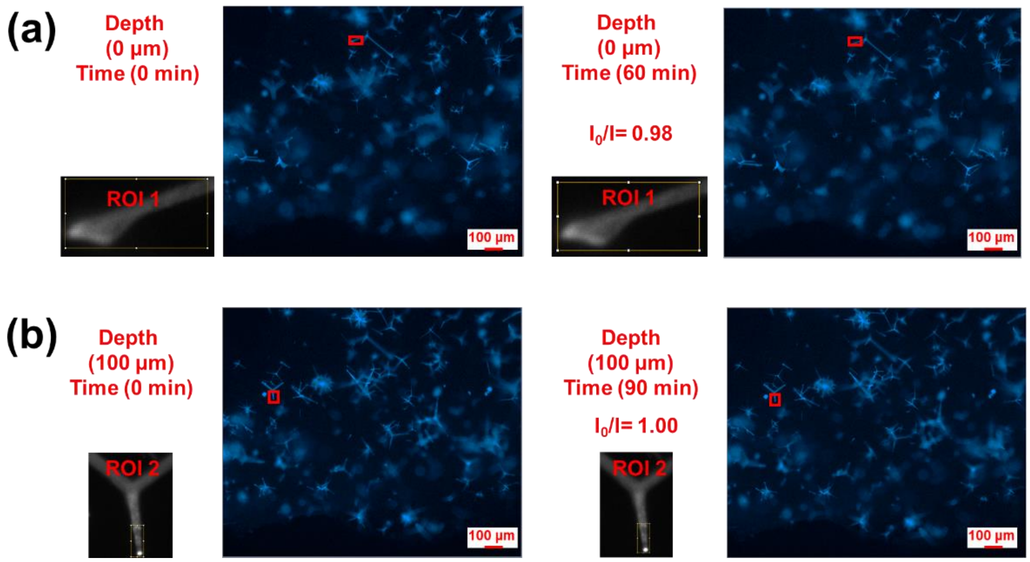


**Figure S10.** 3D t-ZnO sensing without cells. 3D dopamine sensing at different depths of 3D printed construct and time point (a) 0 µm and 60 min, (b) 100 µm and 90 min.

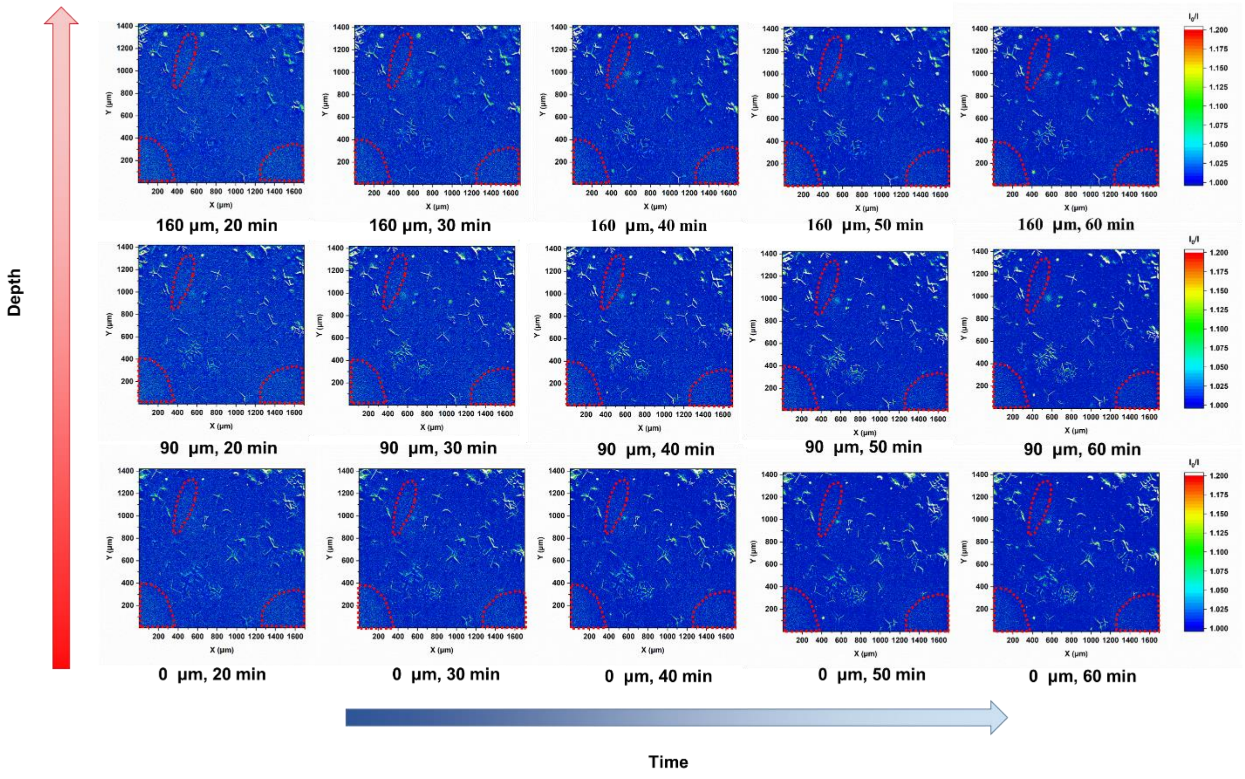


**Figure S11.** Intuitive depiction of 3D DA activity in 3D printed neural tissue after 7 days culture at different imaging time points and structural depths. 3D printed channels are identified with dashed red selections.

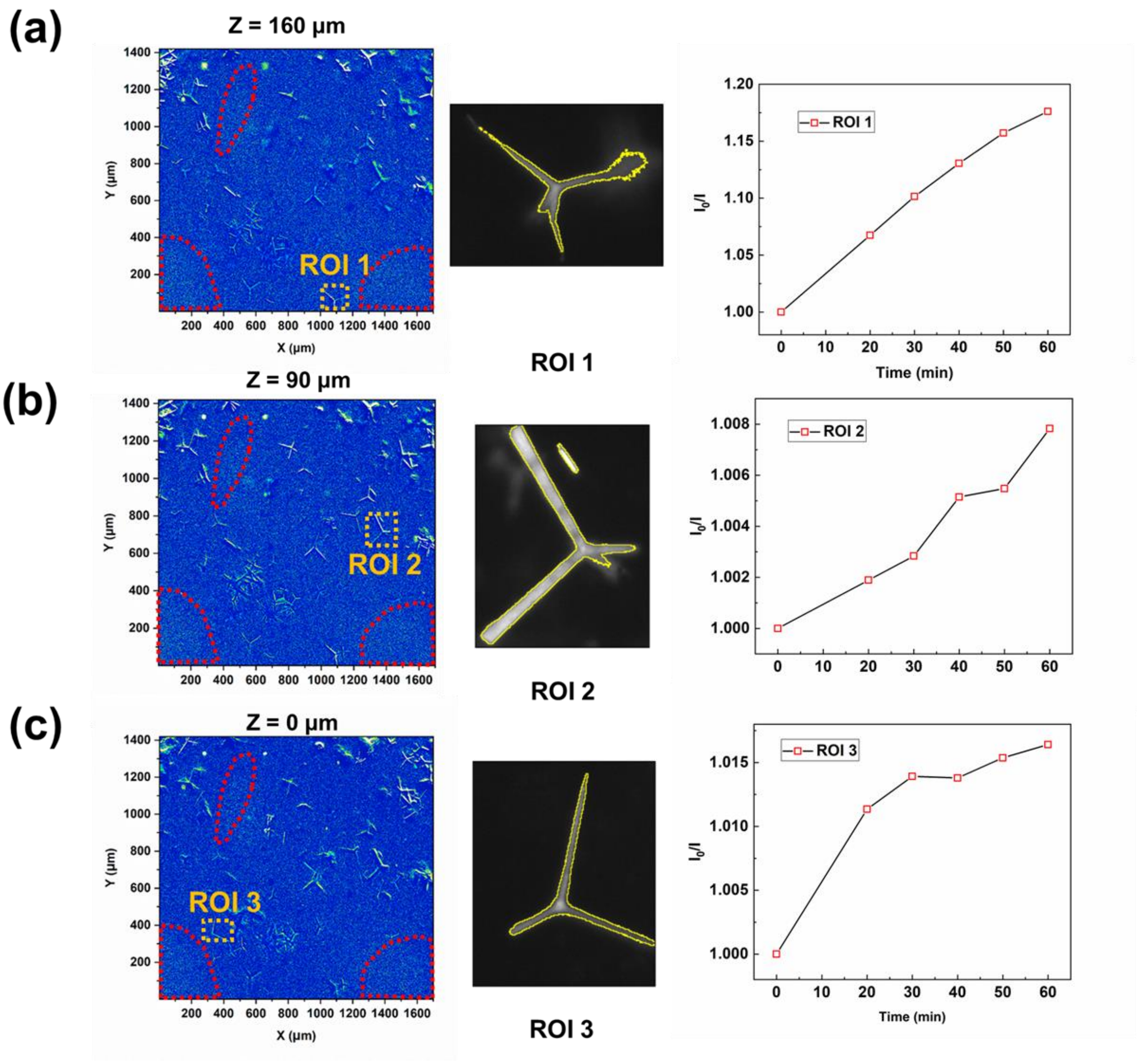


**Figure S12.** t-ZnO based DA sensor in 3D printed tissue. Three ROIs at different depths of (a) 160 µm, (b) 90 µm and (c) 0 µm with corresponding time-dependent quenching curves of t-ZnO emission as $I_0/I$ versus time. t-ZnO structures in the ROIs are highlighted by yellow contours.

**Table S1.** Influence of dilution on the recovery ratio of spiked dopamine in differentiation medium (DM).

| Dilution ratio (PBS:DM) | Spiked dopamine concentration (µM) | Measured concentration (µM) | Recovery of dopamine (%) |
|---|---|---|---|
| 0:1 | 300 | 198 ± 2.2 | 66 |
| 9:1 | 300 | 321 ± 4.8 | 107 |
| 1:0 | 300 | 319 ± 3.3 | 106 |

**Table S2.** Detection of dopamine in diluted DM (PBS:MD = 9:1).

| Spiked dopamine concentration (µM) | Measured concentration (µM) | Recovery of dopamine (%) |
|---|---|---|
| 600 | 629 ± 19.2 | 105 |
| 200 | 209 ± 2.1 | 104 |
| 50 | 53 ± 1.0 | 106 |

**Video S1.** 3D printing of 2%/30%/5% (w/w) Alg/Plu/t-ZnO composite.

**Video S2.** Quenching of t-ZnO autofluorescence after addition of 1 mM dopamine. Recording were performed on a Zeiss Axio Observer microscope.
